# Remodeling of *Il4-Il13-Il5* locus underlies selective gene expression

**DOI:** 10.1101/2022.07.22.501024

**Authors:** Hiroyuki Nagashima, Franziska Petermann, Aleksandra Pękowska, Vijender Chaitankar, Yuka Kanno, John J. O’Shea

**Affiliations:** Lymphocyte Cell Biology Section, NIAMS, NIH, Bethesda, MD 20892, USA; Laboratory of Lymphocyte Nuclear Biology, NIAMS, NIH, Bethesda, MD 20892, USA; Dioscuri Centre for Chromatin Biology and Epigenomics, Nencki Institute of Experimental Biology, Polish Academy of Sciences, 3 Pasteur Street, 02-093 Warsaw, Poland

**Author notes:** Correspondence: J.O’S.

## Abstract

The type 2 cytokines, interleukin (IL)-4, IL-5 and IL-13 reside within a tandem multi-gene cluster in mammals. These cytokines represent the hallmark of type 2 immune responses controlling parasites, promoting tissue repair as well as causing allergic diseases. Both innate and adaptive lymphocytes secrete type 2 cytokines with discordant production spectra. We took a holistic structural and functional view of the type 2 cytokine locus before and after activation, comparing innate (ILC2) and adaptive (Th2) lymphocytes to understand mechanisms underlying their distinctive programs. Rapid induction of IL-5 dominates in ILC2, whereas IL-4 does so in Th2 cells. Using high-resolution chromatin conformation capture we found that global cellular chromatin architecture remained constant, whereas the type 2 cytokine locus rapidly remodeled. In ILC2, *Il13* and *Il5* loci were aligned in proximity whereas *Il4* locus was insulated. In Th2 cells, *Il4* and *Il13* positioned in proximity while the *Il5* locus remained distal. Select REs were separately deleted in mice to confirm cell-type specific and activation-dependent roles in type 2 responses *in vivo*. Thus, contrary to the premise that chromatin architecture plays a minimal role in steady-state gene induction, signal-dependent remodeling of 3D configuration underlies the discordant cytokine outputs in ILC2s versus Th2 cells.

## Main text

Allergy and chronic inflammation are a worldwide health problem affecting billions of peoples and the rise in prevalence of allergic diseases has continued for the past decades. Multiple factors both genetic and environmental influence prevalence and severity of allergic responses. One of the core culprits driving pathological allergic responses is dysregulated production of type 2 cytokines by specialized cytokine-producing lymphocytes. There are two arms of developmentally distinct lymphocytes serving as major producers of type 2 effector cytokines: innate lymphoid cells (ILC2) and T helper cells (Th2)^1, 2^. Balanced and controlled action from ILC2 and Th2 is critical for host defense, barrier integrity, tissue repair and lipid metabolism in health whereas dysregulation leads to various forms of allergic diseases^3–6^. Delineating shared and unique attributes of ILC2 and Th2 are important to understand how dynamic and distinctive type 2 responses are orchestrated by the immediate action of ILC2 and the delayed stepwise Th2 differentiation. Multiple effector cytokines including IL-4, IL-5, IL-6, IL-9, IL-13, granulocyte/macrophage colony stimulating factor (GM-CSF) and amphiregulin comprise type 2 responses, among which IL-4, IL-13 and IL-5 constitute a unique gene cluster in *cis*, *Il4-Il13-Il5*, indicative of coordinated transcriptional regulation of the three cytokines^7–9^. Despite their similarities and shared modes of regulation, ILC2s are the major source of IL-5, which drives eosinophilic disease. By contrast, Th2 cells produce IL-4 whereas ILC2s cells do not. Only with further differentiation do Th2 cells acquire the capacity to make high levels of IL-5^6^.

Elucidation of the mechanisms underlying the selective cytokine production in two related cells provides new insights into gene regulation. Interestingly, this genomic cluster also includes a locus control region (LCR) first identified in mice^10^ and conserved in humans, indicative of potential specialized intrinsic regulatory mechanisms controlling the extended loci. Genome-wide association studies (GWAS) have revealed single nucleotide polymorphisms (SNPs) residing in proximity to the *Il4-Il13-Il5* linked to allergic disease risk^11^. However, there is still a knowledge gap between disease manifestation and genetic variations and the precise molecular mechanisms controlling selective cytokine expression. While multiple layers of regulatory mechanisms are expected in play including transcription factor (TF) loading, chromatin accessibility or higher order chromatin architecture, a detailed understanding of contribution from those regulatory mechanisms remain elusive. Here, we sought to take a holistic view of type 2 cytokine loci before and after activation, comparing ILC2s and Th2 cells, with a particular focus on the interaction between 3D genome and TFs, to understand mechanisms underlying cytokine regulation.

### Genome-wide gene expression, TF distribution and chromatin architecture in ILC2s versus Th2 cells during homeostasis and following activation

ILC2s are abundant in mucosal tissues and provide the first line of defense against microbial insults. ILC2s lack adaptive antigen receptors, instead, they produce type 2 cytokines in response to cytokines, neuropeptides, neurotransmitters, hormone, and lipid metabolites derived from tissue-microenvironment in which ILC2s reside^4, 5^. In the lung, IL-33, an IL-1 family cytokine that activates NF-κB and AP-1 pathways, is a primary activator of ILC2s. The neuropeptide neuromedin U (NMU) augments ILC2 function in concert with IL-33 through the activation of NFATs^12–14^.

In response to a short (4 hr) exposure to IL-33 and NMU, 1064 genes out of 20667 protein-coding genes (CCDS Database) were upregulated in ILC2s whereas 902 genes were induced in Th2 cells upon crosslinking of the T cell receptor (TCR) and the costimulatory molecule CD28 (**Fig. 1a and b, Table 1**). Upregulated genes were classified based on degree of induction and designated as weakly inducible genes (WIGs, >10 FPKM and <100 FPKM) and highly inducible genes (HIGs, >100 FPKM). *Il5* and *Il13* were by far the highest HIGs in ILC2s, whereas *Penk, Ccl1, Il13, Il10, Il4*, and, to a lesser extent *Il5*, were highly induced in Th2 cells. Venn diagrams depict the common or cell-type specific HIGs and WIGs (**Fig. 1c, Table 2**). There were 45 common HIGs shared with ILC2s (out of 115 HIGs) and Th2 (out of 94 HIGs), which included *Ccl1, Il13, Irf4, Csf2, Il2, Batf, Il5* and *Myc*, while 49 Th2-specific HIGs included *Il3, Il4, Il10* and *Areg*. These data are consistent with previous studies showing the absence of IL-4 production in ILC2s^14^. *Areg*, encoding amphiregulin, was not induced in ILC2s with this stimulation condition as we reported previously^14^.

**Figure 1.**
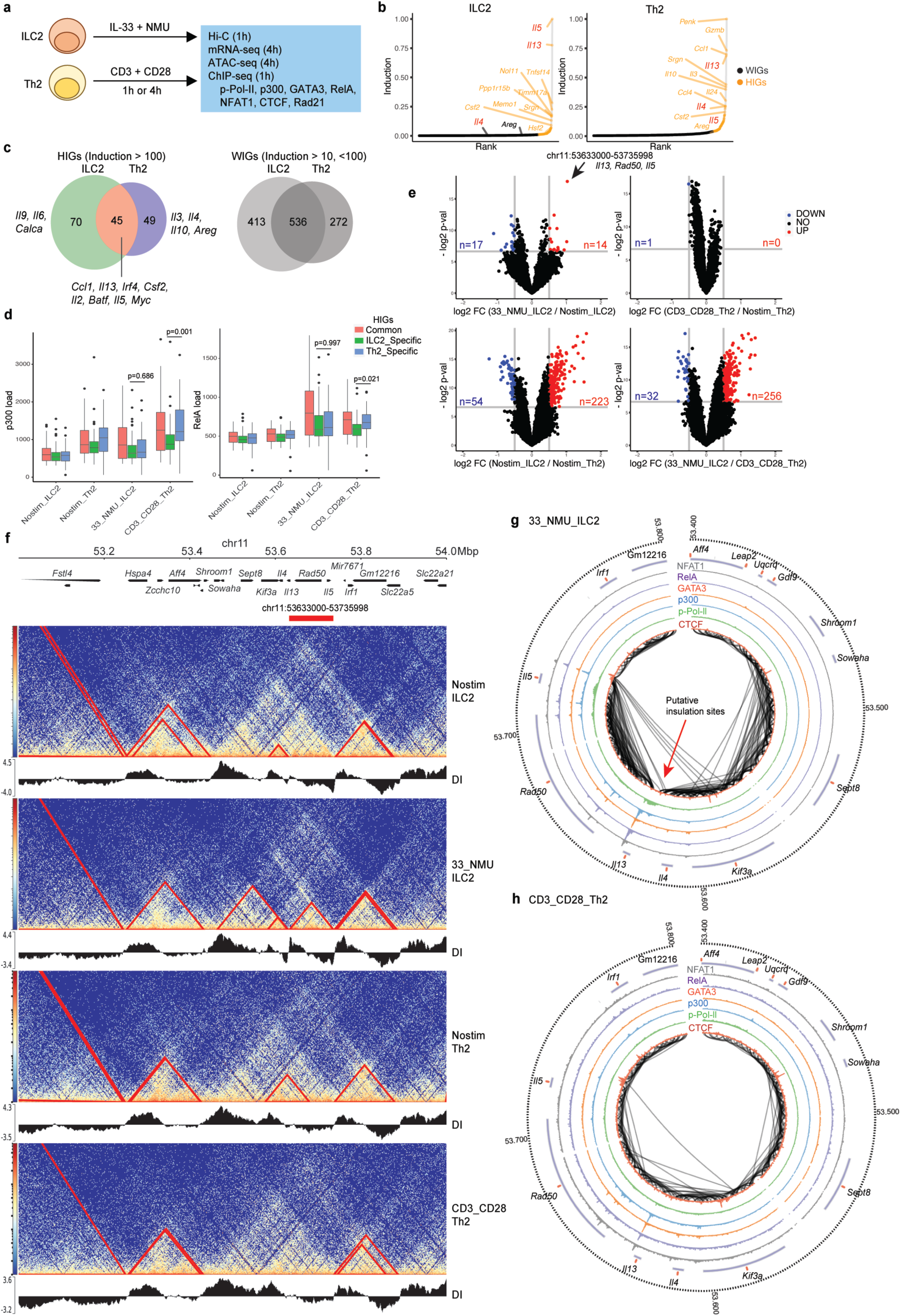
Genome-wide gene expression, TF distribution and chromatin architecture in ILC2s and Th2 cells during homeostasis and following activation. **a**, Experimental design for *in situ* Hi-C, mRNA-seq, ATAC-seq and ChIP-seq using *in vitro* expanded ILC2s and Th2 cells. ILC2 data for mRNA-seq and ATAC-seq were previously reported (GEO) **b**, Magnitude of gene induction, confirmed by mRNA-seq (FPKM (Stim - Nostim)). The genes with an increase of more than 10 FPKM upon stimulation were plotted and classified into weakly inducible genes (WIGs, < 100, > 10 FPKM) and highly inducible genes (HIGs, > 100 FPKM) based on gene induction. The Y axis represents normalized gene induction. **c**, Venn diagram depicts common and specific HIGs and WIGs between ILC2s and Th2. **d**, Aggregated TF loads near HIGs (+/-50kb from TSS) were calculated and binned into 3 categories (common-, ILC2 specific- or Th2 specific) and graphed (Y-axis) for 4 sample conditions (X-axis). p300 loads (left panel), RelA loads (right panel). **e**, For a total of 20857 topologically associated domains (TADs) identified, pair-wise comparisons were performed to select for significantly different TADs (red/blue colored) due to stimulation (top panels) or cell type (bottom panels). **f**, HiC heatmaps for 4 samples in a window of 53.1 - 54 Mbp on chromosome 11. DI (directionality index) track accompanies each heatmap at the bottom. The position of the TAD specific to activated ILC2 is marked by a red bar at the top. **g, h**, Circus plots of the extended type 2 cytokine loci depicting HiC interactions in stimulated ILC2s (top) and stimulated Th2 (bottom). ChIP-seq based TF footprints are also shown as multiple circles for NFAT, RelA, GATA3, p300, Pol II and CTCF. A representative data from two or more biological replicates were used to graph each component.

ILC2-specific HIGs comprised 70 genes including *Il9* and *Calca* encoding IL-9 and CGRP respectively^14^. To identify potential molecular mechanisms controlling cell-type specific HIGs, we first quantified the aggregate enrichment of the histone acetyltransferase (HAT), p300 and the NF-κB subunit, RelA, over the common or specific HIGs using ChIP-seq (**Fig. 1d**) considering that a sum of HAT or TF loading could be a major driver and predictor of gene induction. In Th2 cells, increased p300 and RelA loads both positively correlated with Th2-specific HIGs (blue bars) and commonly induced HIGs (red bars). However, in ILC2 cells there was no difference between TF and HAT loads between ILC2-specific HIGs (green bars) and Th2-specific HIGs. This implied a simple model for HIGs regulation in Th2 cells – TF and HAT load correlate directly with magnitude of gene induction. On the contrary for ILC2s, ILC2-specific HIGs were induced through a mechanism other than recruiting more transactivating factors (**Fig. 1d**).

Based on these results, we next hypothesized that the higher-order chromatin architecture might contribute to gene regulation and serve as a mechanism to confer cell-type specific induction of HIGs in ILC2. Higher-order chromatin architecture comprises A (“active”) and B (“inactive”) compartments, topologically associating domains (TADs) and promoter–enhancer interactions to provide a fundamental landscape for gene regulation^15, 16^. Active involvement of 3D genome structure is implicated during stepwise progress of cellular development for cell fate decision and function^17–20^. After a cell is committed to a cell state, the three-dimensional chromatin regulated by cohesin has been reported to not be required for proper gene expression^21, 22^. However, cohesin is required for inducible enhancer activity that underlies inflammatory gene expression in macrophages upon response to microbial signals^23^. Therefore, we hypothesized that chromatin architecture underlies differential stimulation-dependent gene regulation between ILC2s and Th2. To test this hypothesis, we performed *in situ* Hi-C using ILC2s and Th2 cells and identified a total of 20857 topologically associated domains (TADs) that were relevant to at least one cell-type or stimulation condition tested (**Fig. 1a and e, Extended Data Fig. 1, Table 3**). Pair-wise comparisons of TADs between 2 conditions (homeostasis vs activation or ILC2 vs Th2) revealed that intrachromosomal interactions differed more between cell types than between activation status (**Fig. 1e, Table 3**). While the short activation window (1 hr) did not impact overall genome conformation in Th2 cells or ILC2s, a segment of the type 2 cytokine locus (chr11: 53633000 - 53735998) was a notable exception in ILC2s, as it rapidly remodeled following cell activation (**Fig. 1e: top left, Fig.1f: red bar**). The extended type 2 cytokine locus ranging from *Shroom1* to *Il5* formed a *de novo* TAD composed of *Il13, Rad50* and *Il5* upon acute stimulation in ILC2s, and the contact frequencies were notably higher than in Th2 (**Fig. 1f-h**). Furthermore, there was a putative insulator region identified only in ILC2s (**Fig.1g, red arrow**). This suggested that the cell-type specific chromatin conformation correlated with the ability of ILC2s to generate massive amounts of *Il5* and *Il13*, but not *Il4* (**Fig.1b**).

We confirmed that other effector cytokine loci, *Il9* (ILC2 specific) and *Csf2*-*Il3* (Common and Th2 specific, respectively), also showed cell-type specific and activation dependent conformational changes, consistent with their gene expression profiles (**Fig. 1c, Extended Data Fig. 2 and 3**).

Interchromosomal interactions between the extended *Il4/Il13/Il5* and *Ifng* loci have been reported^24^. The *Ifng* gene reportedly interacts with the *Il5* promoter, *Rad50* promoter and LCR. Deletion of the Th2 LCR affects not only production of type 2 cytokines in Th2 but also IFN-γ production in Th1. Therefore, we sought to access global interchromosomal interactions comparing ILC2 and Th2 cells. Although signals were significantly weaker in comparison to intrachromosomal interactions, interactions across distinct chromosomes were detected (**Extended Data Fig. 4 and 5**). Specifically, the interaction between chr11:53MB -55.5MB containing *Il4/Il13/Il5* as well as *Csf2/Il3*, and chr10:117MB - 118.5MB containing *Il22/Ifng* were detected in ILC2s (**Extended Data Fig. 5a**). This broad interaction diminished following stimulation with IL-33 plus NMU, suggesting remodeling of interchromosomal interaction during activation (**Extended Data Fig. 5b-c**). Since the percentage of interchromosomal interaction was low (less than 20%, **Extended Data Fig. 1c**), further high-resolution analysis to address the interchromosomal interaction between *Il4/Il13/Il5* and *Ifng* was not possible.

### Re-defining regulatory DNA elements (REs) controlling type 2 cytokine locus

Previous studies have characterized multiple REs in the vicinity of *Il4/Il13/Il5* loci using DNase I hypersensitive assays or DNA sequence comparison across species. These include: the Th2 LCR composed of DNase-I hypersensitive site (HS) of *Rad50* (RHS)-4, -5, -6 and -7, the conserved GATA3-responsive element (CGRE), conserved noncoding sequence 1 (CNS1), HS-II, HS-IV and HS-V^7–9, 25^; functional validation of these loci was accomplished by deletion of the endogenous locus in mice. We identified additional REs controlling this locus by integrating the chromatin structure, gene accessibility and TF footprint (**Fig. 2**: red fonts). Those novel REs were classified into two groups, structural and functional. The former (structural REs) were defined by binding of either or both DNA-binding protein CCCTC-binding factor (CTCF, blue) and Rad21 (orange), a component of the cohesin complex, which cooperatively establish chromatin loops and also form insulation sites^16, 21, 26^. Functional REs were defined by binding of p300 (yellow), a lineage-determining TF (LDTF) (e.g. GATA3, black) or stimulation-dependent TFs (SDTFs) including RelA (green) and NFAT1 (data not shown). The structural REs included the following potential elements: *Shroom1*-associated SHS-I and SHS-II, *Kif3a*-associated KHS-I and KHS-II, and *Il5*-associated 5HS-IIIa, b and c. Interestingly, an element denoted as +6.5kb*^Il13^* located 6.5kb downstream from *Il13* promoter and 1kb upstream from CNS-I was a putative insulator bound to CTCF and Rad 21, and formed a boundary between *Il4* and *Il13* genes (**Fig. 1f-g** **and** **Fig. 2**). The functional enhancers included 5HS-Ia, b, c, d and e, and 5HS-II, which are in proximity to *Il5*. Among them, +6.5kb*^Il13^* and 5HS-Ib were found to be evolutionary conserved (phyloP Score), suggesting their important roles for controlling type 2 cytokines.

**Figure 2.**
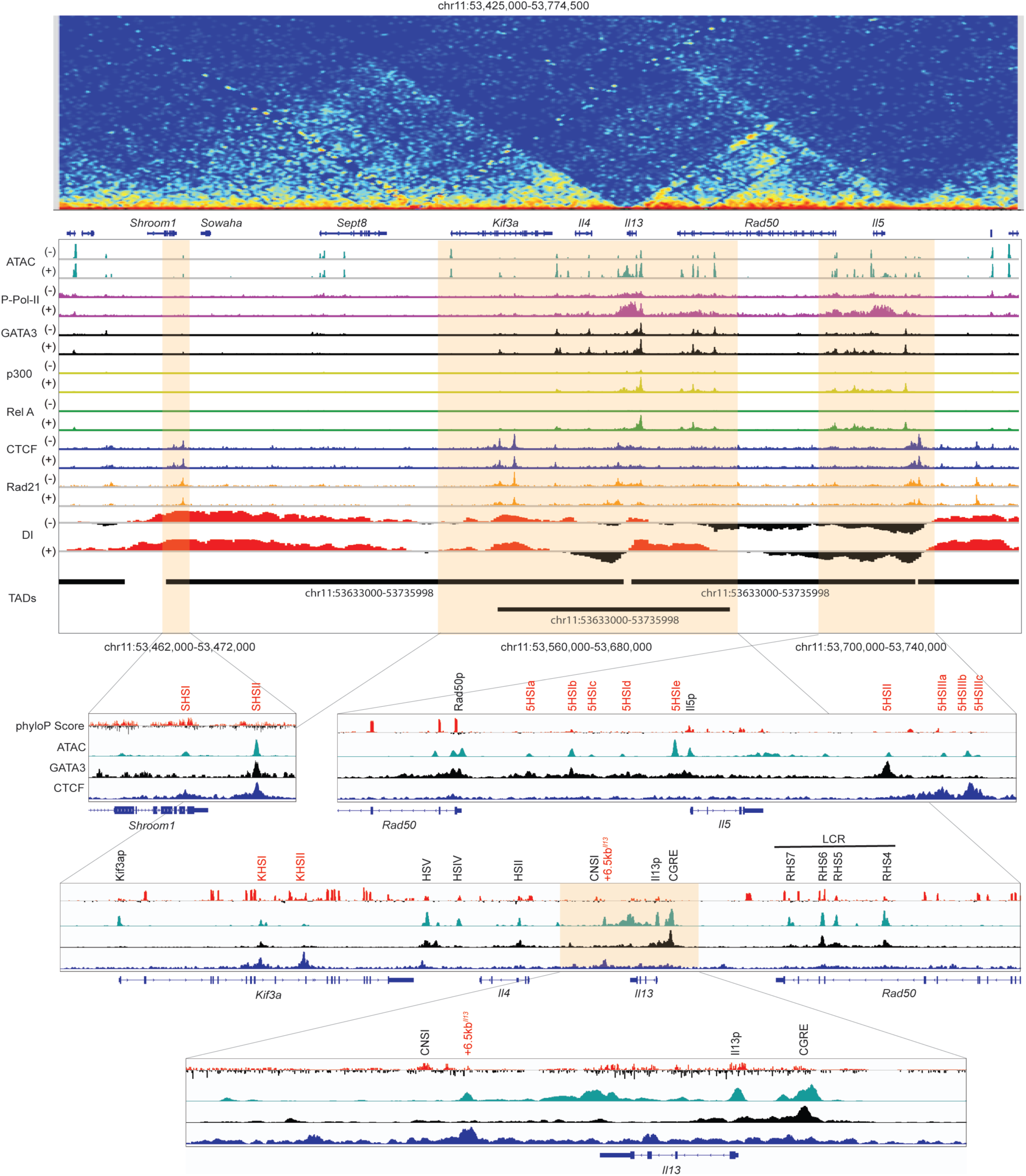
Re-defining REs controlling type 2 cytokine locus. Integrated multi-layered genome track view of type 2 cytokine locus (53,425,000-53,774,500 on chromosome 11) in ILC2. From the top, HiC heatmap, ATAC-seq, ChIP-seq for 6 factors indicated, DI, phyloP Score (Mammal Basewise Conservation of 59 vertebrate) and TADs. ATAC and ChIP were performed using ILC2s without (-) or with (+) stimulation to see activation induced changes. Four select regions in orange shade were further magnified and shown below, in which known (black fonts) and new (red fonts) regulatory regions are annotated based on ATAC-seq peaks. Representative data from two biological replicates were used for presentation.

### Interplay among TFs and type 2 cytokine locus in 3D space

To better understand how the interaction between TFs and REs control chromatin remodeling, we modeled 3D structure of type 2 cytokine locus using GenomeFlow^27^ based on Hi-C data (**Fig. 3**). ILC2 stimulation with IL-33 in combination with NMU brought CGRE, LCR (RHS(5+6)) and 5HSI(a-d) together in proximity, and also juxtaposed +6.5kb*^Il13^* and 5HS-III(a-c) (**Fig.3a, b, e and g**). In contrast, Th2 stimulation with TCR and CD28 led to more REs being brought into vicinity including HS-II, +6.5kb*^Il13^*, KHS-I, HS-V, CGRE and LCR **(Fig.3c, d, f and g).** This correlated with a stimulation-dependent and a cell-type specific recruitment of p300, RelA and NFAT1, indicating that SDTFs loading associates with chromatin remodeling and juxtaposition of enhancers (**Fig. 1g-h, 3 and Extended Data Fig. 6**). The interaction between CGRE and LCR (RHS(5+6)) was stable and unchanged with activation in both cells (**Fig.3e-f)**, suggesting that this interaction had already been established prior to stimulation (**Fig.3g)**, presumably via constitutive GATA3 binding (**Fig. 2**). This observation is in line with previous reports describing an indispensable “structural” role of LCR in controlling the long-range interaction of type 2 cytokine locus and their expression^10, 28–32^.

**Figure 3.**
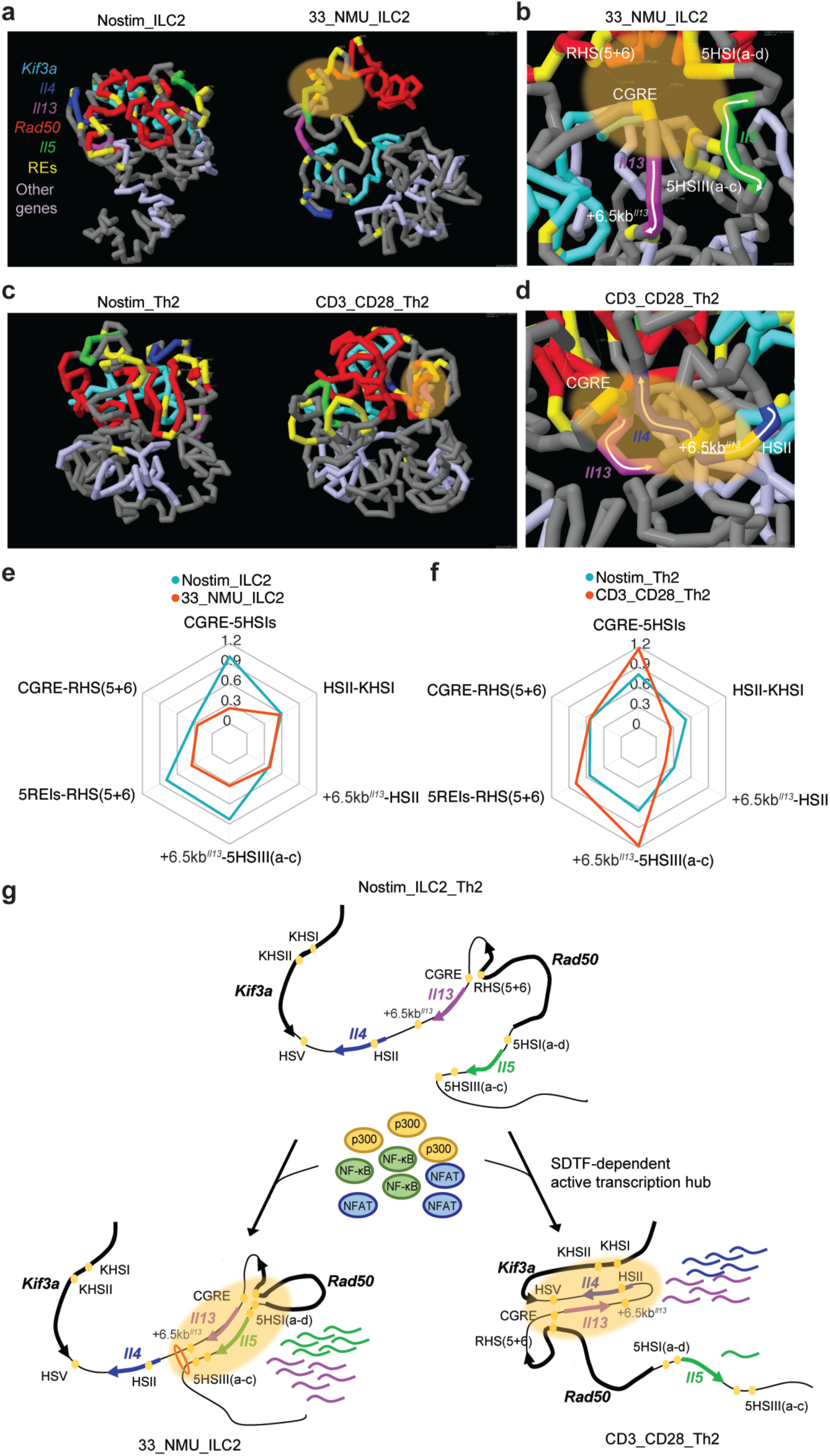
Interplay among TFs and type 2 cytokine locus in 3D space. **a-d**, 3D models of type 2 cytokine locus generated by GenomeFlow using HiC interaction data from the region spanning from 53,425,000 to 53,775,000 (350KB) on chromosome 11. ILC2 (**a, b**) and Th2 (**c, d**). In the magnified views (b, d), REs are colored in yellow and annotated. White arrows depict directions of mRNA transcription for cytokine genes. **e, f**, Radar plots showing distance between two REs in no stimulation (blue) or stimulation (red) for ILC2 (**e**) and Th2 (**f**). Distances were calculated by GenomeFlow with HiC data. Y axis represents nanometer. **g**, 2D presentation of topology among REs and 3 cytokine genes in non-activated cells (top), activated ILC2 (bottom left) and activated Th2 (bottom right), leading to discordant cytokine productions between the 2 cell types.

The loop extrusion of chromatin controlled by cohesin is thought to promote genome segregation and formation of the *de novo* boundaries to control gene expression. In this model, cohesin binds to chromatin, reels it in, and extrudes it as a loop^33–35^. To understand the contribution of trans factors controlling a *de novo* boundary formed by interacting +6.5kb*^Il13^*and 5HS-III(a+b+c) in ILC2s, we examined Rad21 binding by ChIP-seq. We found that Rad21 constitutively bound to the structural REs, namely +6.5kb*^Il13^*, 5HS-III(a+b+c), KHS-I and KHS-II, and this binding was constant regardless of cell activation status (**Fig. 2: orange tracks**). Thus, it is unlikely that loop extrusion occurred after ILC2 stimulation, at least in this short time window of stimulation. Instead, the binding of SDTFs such as NFκB-RelA (**Fig.2: green tracks**) to select inducible enhancers marked by p300 (**Fig.2: yellow tracks**) appeared to be a major contributor driving topological structural alteration. Collectively, we identified multiple novel REs that employed select targets of different classes of TFs including SDTFs and were interspersed among structural REs heavily bound by CTCF/Rad21. Precise positioning of structural and functional REs presumably allows TF network to coordinately control chromatin remodeling and gene expression in a cell-type specific manner.

### Critical role of enhancers and structural REs during allergic lung inflammation

To experimentally validate predicted functional roles of REs identified, we next generated targeted deletion of 6 REs in mice: KHS-I, KHS-II, 5HS-I(a-d), 5HS-Ie, 5HS-II and 5HS-IIIs-(a-c) respectively (**Fig. 4a** **and table 4**). Homozygous deletion of KHS-II (chr11:53590014 – 53591069) in mice resulted in embryonic lethality, likely due to the loss of exon 10 in *Kif3a* which is important for early embryonic development^36^ (data not shown). Thus, we tested the remaining five strains. Intranasal administration of the cysteine proteinase, papain, induced chronic lung inflammation along with strong activation of ILC2s and Th2 cells in wild-type (WT) mice (**Fig. 4b-f**).

**Figure 4.**
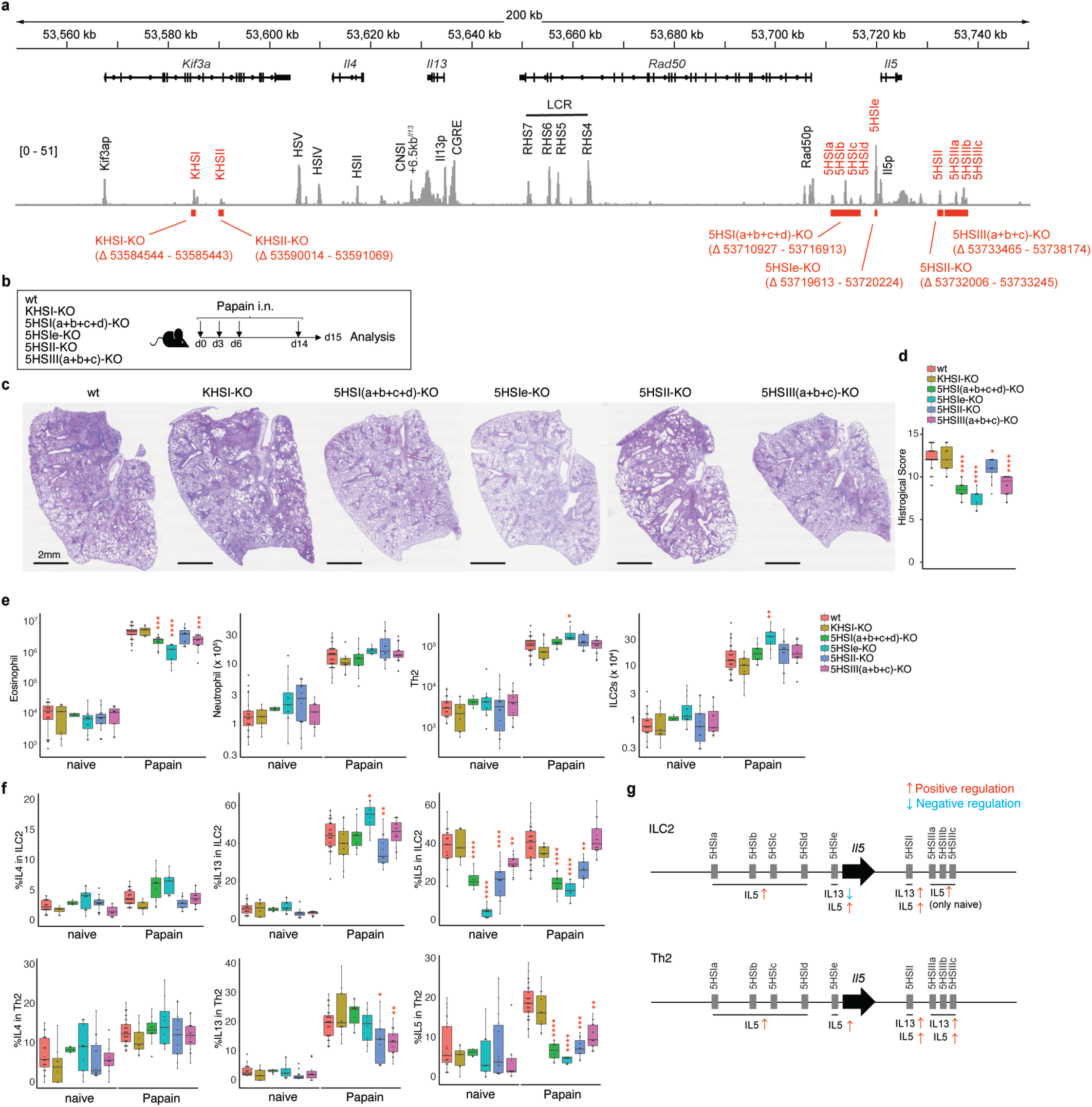
Critical role of enhancers and structural REs during allergic lung inflammation. **a**, REs identified in this study were selectively deleted (red bars) to generate 6 mouse strains, and subjected to papain induced lung inflammation (**b**) to compare their type 2 response. WT (naïve; n=24, papain; n=37), KHS-KO (naïve; n=8, papain; n=11), 5HS-I(a+b+c+d)-KO (naïve; n=4, papain; n=10), 5HS-Ie KO (naïve; n=12, papain; n=8), 5HS-II KO (naïve; n=14, papain; n=11), and 5HS-III(a+b+c) KO (naïve; n=11, papain; n=16) were analyzed in two to five independent experiments. **c**, PAS staining of the lung from mice treated with papain. A representative image from each genotype is shown. **d**, Histological scores were calculated for each genotype group and compared with statistical evaluation (asterisks). **e**, Cell numbers recovered from the lungs were compared among 6 genotypes (color code as in d) in naïve and papain challenged conditions. Cells evaluated were eosinophils (CD11b^+^ Siglec-F^+^), neutrophils (CD11b^+^ Gr1^+^), Th2 (CD3ε^+^ TCRβ^+^ CD4^+^ CD44^+^ Foxp3^-^ GATA3^+^) and ILC2s (Lin^-^ Thy1^+^ CD127^+^ GATA3^+^). **f**, Cytokine production from ILC2s (top panels) and Th2 (bottom panels) were evaluated by FACS for IL-4, IL-13 and IL-5. **g**, Graphical summary of impact on cytokine production due to RE deletion in mice. Cell type dependent differential impact was noted for some REs. Statistical significance is depicted as ∗p < 0.05, ∗∗p < 0.01, ∗∗∗p < 0.001 and ∗∗∗∗p < 0.0001 (Student’s t test).

Compared with WT responses, we observed less lung inflammation with decreased eosinophil infiltration, minimal goblet cell hyperplasia, and reduced mucus production in 5HS-I(a+b+c+d) KO, 5HS-Ie KO, and 5HS-III(a+b+c) KO mice, evaluated by aggregated histological scores (**Fig, 4d**). In 5HS-II KO mice, the alternations in these inflammation parameters were less evident and KHS-I KO mice exhibited comparable responses to WT (**Fig. 4c-e**). The number of neutrophils, Th2 and ILC2s were largely comparable except for a modest increase in Th2 and ILC2s in papain-treated 5HS-Ie KO mice. Loss of each RE resulted in varied impact on type 2 cytokine expression in ILC2s and Th2 cells, among which the most striking effect being the lack of IL-5 production in 5HS-Ie KO mice (**Fig.4f**) that correlated with selective reduction in eosinophils (**Fig. 4e**) while sparing IL-13 production. Thus, 5HS-Ie acted as an essential enhancer for IL-5 expression in both ILC2s and Th2 during homeostasis and inflammation.

A similar phenotype was seen for 5HS-I(a-d), with impaired IL-5 production in these mice being associated with reduced eosinophil response following papain challenge (**Fig. 4e**). By contrast, loss of 5HS-II (blue) impaired IL-5 and IL-13 equally in ILC2s and Th2 cells, suggesting its role as both *cis-* and *trans*-acting enhancer (**Fig. 4f and g**). Compared with other REs, the impact of 5HS-III(a+b+c) deletion (purple) was most context dependent, exhibiting a cell-type specific and condition-dependent function. For instance, IL-5 expression of ILC2s from 5HS-III(a+b+c) KO mice prior to papain treatment was decreased compared to that of WT ILC2s. By contrast, no difference in IL-5 expression was seen in ILC2s from WT or KO mice following papain treatment. Also, loss of 5HS-III(a+b+c) led to defective IL-13 and IL-5 production in Th2 cells during inflammation, but had no effect on cytokine production in ILC2s. Moreover, comparable amounts of papain-specific IgM, IgG1 and IgE were detected in serum of 5HS-Ie KO, 5HS-II KO and 5HS-III(a+b+c) KO mice, suggesting that humoral response and immunoglobulin class switching in B cells were maintained that were driven by follicular Th (Tfh) cells producing IL-4 (data not shown).

Collectively, the phenotype from RE deleted mice indicated that ILC2s and Th2 cells utilize the distinct REs to control type 2 cytokines during allergic reaction. Notably, the observed *in vivo* phenotype in RE-targeted mice was in line with our earlier observation with *in vitro* stimulated cells in which 3D configuration of type 2 cytokine locus differs between ILC2 and Th2 cells (**Fig. 3g and h**).

### Structural REs downstream of Il5 is critical for segmentation of genome of the extended type 2 cytokine locus

Based on the response of the RE mutants, we next considered that a *de novo* boundary might form at +6.5kb*^Il13^* upon activation and together with 5HS-III(a-c) functions as an insulator. (**Fig. 3g**). Using cultured ILC2s stimulated *in vitro*, we confirmed that 5HS-III(a-c) deletion exaggerated IL-4 mRNA production while limiting IL-5 mRNA production (**Fig. 5a, b**).

**Figure 5.**
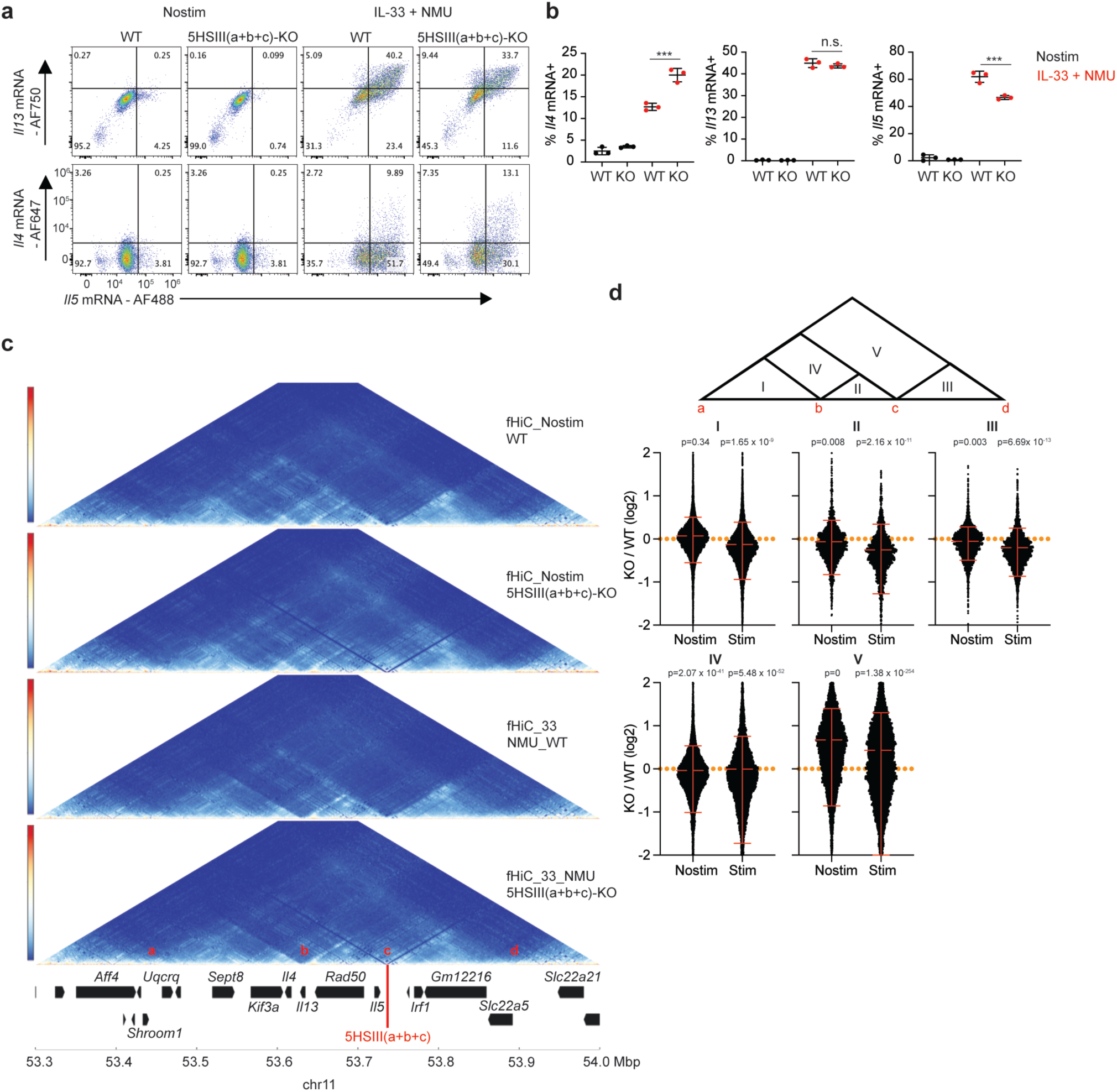
Structural REs downstream of Il5 is critical for segmentation of genome of the extended type 2 cytokine locus. **a, b**, Type 2 cytokine mRNA expression detected by PrimeFlow in ILC2s stimulated with IL-33 plus NMU stimulation. WT and 5HS-III(a+b+c) KO ILC2s were compared. Shown in **a** were representative plots from three independent experiments, and triplicates summary were shown in **b**. **c**, HiC heatmaps depicting intrachromatin interactions within type 2 cytokine locus in ILC2 using focused Hi-C (fHiC) method. Conditions tested were WT – no stimulation, 5HS-III(a+b+c) KO – no stimulation, WT – stimulation (IL-33 + NMU) and KO – stimulation. **d**, The quantification of HiC interaction within five segments shown at the top. For each segment, frequencies of interactions were quantitated, and plotted as KO/WT ratio, and any deviation from level 0 (orange line) is indicative of differential segmentation between WT and KO.

To test the possible insulator function of 5HS-IIIs-(a-c), we employed focused Hi-C to capture chromosomal interactions within 1Mbp region containing the type 2 cytokine locus, comparing WT and 5HS-IIIs-(a-c) KO (**Fig. 5c** **and Table 5**). We found that absence of this RE led to a defect in formation of a boundary downstream of *Il5.* This resulted in failure to partition the type 2 cytokine locus from the upstream TAD containing *Irf1*, promoting increased interactions in region V in KO cells (**Fig. 5c and d**, region V). In addition, the intrachromosomal interactions within regions I, II, III in 5HS-IIIs-(a-c) KO ILC2s following stimulation were decreased compared to those of WT ILC2s (**Fig. 5d**). Interestingly, the *de novo* boundary formed at +6.5kb*^Il13^*was intact in KO cells, suggesting that the interaction among SDTFs and enhancers is a key to support this side of *de novo* boundary formation regardless of the condition at 5HS-IIIs- (a-c) end (**Fig. 5c**). Thus, these data demonstrate the requisite role of 5HS-IIIs-(a-c) in localizing chromatin interactions for proper type 2 cytokine responses.

### Relationship between identified REs and human allergic diseases

To evaluate the potential human relevance of our work in mice, we examined publicly available *in-situ* Hi-C and ATAC-seq data of human T cells^37^. In this study, naïve Th cells from peripheral blood were activated with anti-CD3 and anti-CD28 for 3d without additional cytokines. A long-range interaction downstream of *IL5* to *SHROOM1* was evident, as was observed in mice (**Fig. 6**). Additionally, ATAC-seq revealed conserved accessible peaks corresponding to the REs we found in mice. GWAS has revealed a number of SNPs which reside in the type 2 cytokine locus and associate with human allergic diseases such as asthma, atopic dermatitis and rhinitis^11^ (**Fig. 6a**). Of these, three and six SNPs that are likely to reside within 5HS-IIIs and 5HS-Is, respectively. Those SNPs link to the asthma and eosinophil response, consistent with the role of those REs we observed in mice during allergic lung inflammation (**Fig. 6b**). Therefore, the REs we identified in mice are likely to be conserved in human as well and their context dependent regulation of type 2 immunity might be altered by SNPs that modify basal and activation-dependent chromatin organization.

**Figure 6.**
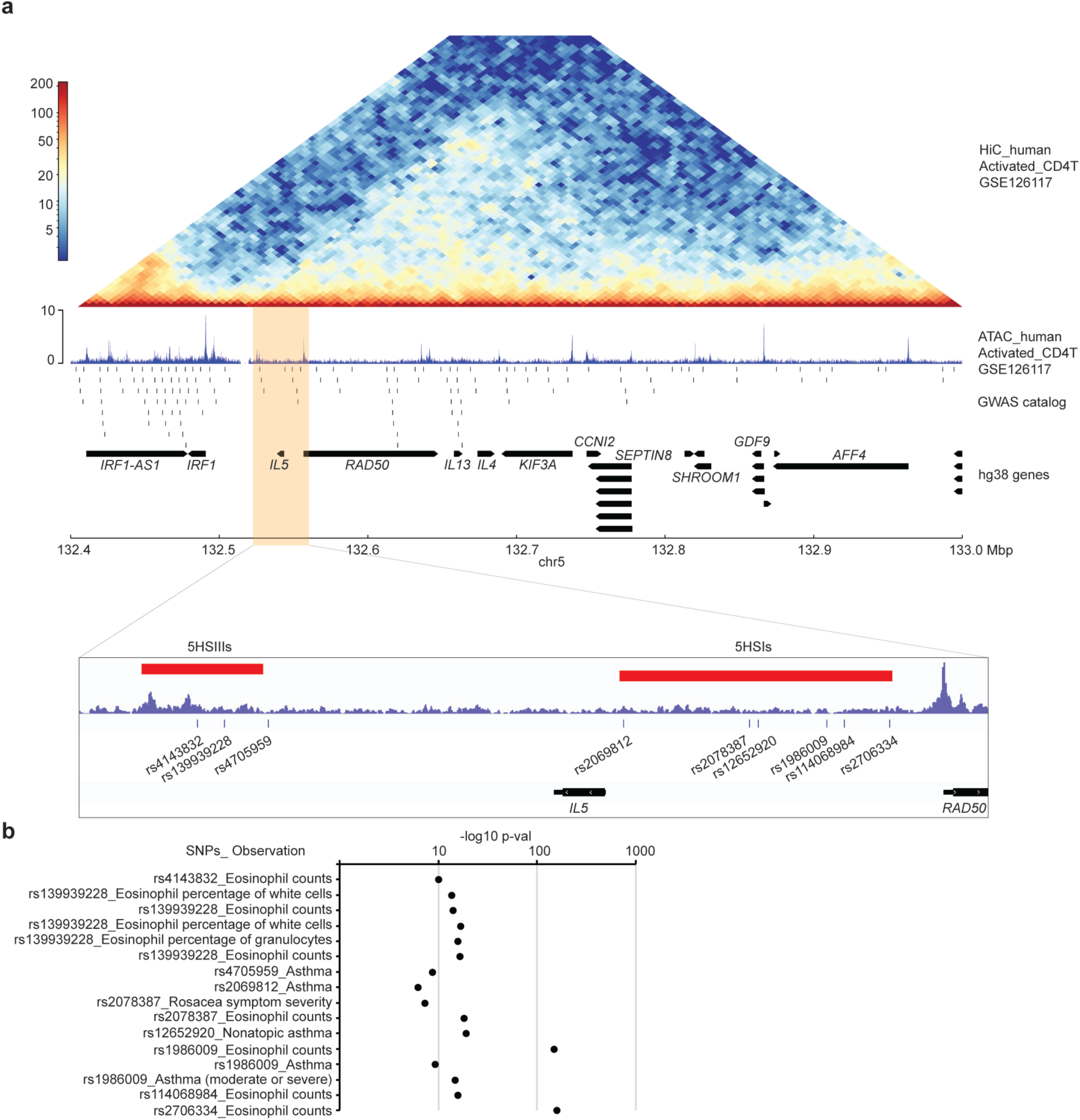
Close connection between REs and SNPs associated with human allergic diseases. **a**, Integration of HiC interaction map, ATAC peaks, and GWAS hits at type 2 cytokine locus. Data were from human Th cells activated with anti-CD3 and anti-CD28 for 3 days available in public database. Predicted regions corresponding to 5HSIs and 5HSIIIs are indicated as red bars. **b**, SNPs that are located within 5HSIs and 5HSIIIs regions and associated with allergy-related traits are listed.

## Discussion

Regulation of the type 2 cytokine locus containing *Il4-Il13-Il5* has attracted intense and continued interest from immunologists and clinicians alike to better understand the basis of coordinated, nuanced and selective gene transcription. Using DNA sequence conservation and enzyme accessibility as research tools, multiple regions across the locus were previously identified and tested for their regulatory functionality in isolation. However, knowledge on comprehensive coordination among putative REs and 3 cytokine loci remained elusive. This study employed an unbiased holistic high resolution (1kb) measurement of chromatin structure to understand topological proximity of REs in 3D space and related this to *trans* factor loading in innate and adaptive lymphocytes, with and without cell activation status and evaluated coordinated functional read outs.

Our findings led us to propose that signal-dependent dynamic 3D configuration instructed by the interaction between TFs and REs is likely to underlie the discordant regulation of type 2 cytokines in ILC2 and Th2 cells in mice. One candidate TF whose expression differs between ILC2 and Th2 is special AT-rich sequence binding protein 1 (SATB1). SATB1 has been reported to regulate the chromatin remodeling of type 2 cytokine locus and the coordinated expression of these cytokines in a mouse Th2 cell line when activated with concanavalin A^38^. We noted that ILC2s expressed much less *Satb1* than Th2 cells (approximately 20 FPKM vs 100 FPKM, data not shown), suggesting that differential SATB1 expression might partly account for the cell-type specific chromatin remodeling of type 2 cytokine locus observed in this study. We also predict that those cell type- and stimulation dependent REs could play roles in context dependent dysregulation of cytokines in human diseases as well.

In response to extracellular soluble stimuli, rapid, massive, and transient induction of the type 2 cytokines ensues, and contrary to the premise that chromatin architecture generally plays a minimal role in steady-state gene induction^21, 22^, dynamic quick remodeling of 3D structure seems like a major underpinning to form cell type specific cytokine production profiles. SDTFs are a key aspect of the machinery for influencing architecture through select functional enhancer type REs embedded in the type 2 locus in ILC2s. We focused on a particular stimulation condition in this study: IL-33 plus NMU for ILC2s, and TCR plus CD28 for Th2 cells, both of which activate NF-κB, AP-1 and NFAT1 pathway. However, this is only a part of the regulation mobilized in these cells. For example, a neuropeptide CGRP counter regulates IL-13 and IL-5 in ILC2s through the activation of cAMP - CREB pathway^14^. Furthermore, type 2 cytokines are produced not only by ILC2s and Th2 cells but also by Tfh, NKT cells, basophils and mast cells. Hi-C experiments give us a snapshot of the aggregate structural view from millions of cells but do not reveal the dynamics of chromatin remodeling. Indeed, a recent report using live cell imaging suggests that chromatin looping is highly dynamic, often in partially extruded states with the fully looped state being rare and transient^39^. Even with population-based assays, our observation is in line with this report revealing the dynamic nature of TADs within the type 2 cytokine locus and their cell type specific preferential configurations. Thus, additional studies will be needed to understand the molecular mechanism by which cell-restricted TFs and distinct SDTFs control state-specific and condition-dependent chromatin remodeling over time. Ultimately obtaining the trajectories of dynamic TAD changes at single-cell levels will be important.

In humans, it is reported that pathogenic Th2 cells producing IL-5 dominate in severe asthmatic lung, and blocking IL-5 is effective in treating asthma patients with severe eosinophilic disease^6^. Our study identified the REs controlling *Il5* and the SNPs associating with eosinophil response, pointing to a possibility of pathogenic Th2 cells carrying variants in *Il5* controlling the function of these REs. In this study, we analyzed publicly available data from non-polarized human T cells. Thus, chromatin remodeling of type 2 cytokine locus in specialized type 2 cells (Th2 and ILC2s) in humans with genetic variation remains elusive. Further analyses with human cells will be needed to understand the evolutionally conserved regulatory machinery for this clinically important gene locus. It is becoming evident that the function of cytokine-producing lymphocytes goes beyond immunity and host defense. Their function also collaborates with body’s metabolism, circadian rhythm, and autonomic function, leading a way to further advance our knowledge on multiple fronts of biology encompassing host defense, tumor immunity, metabolism, stress response, cognitive function and beyond.

## ACKNOWLEDGMENTS

We thank HY. Shih for critical inputs in planning the experiments and analyzing the data. We thank S. Dell’Orso and F. Naz (Genome Analysis Core Facility, NIAMS); J. Simone, J. Lay, and K. Tinsley (Flow Cytometry Section, NIAMS); H-W. Sun, A. Uhlman, K. Jiang and S.R. Brooks (Biodata Mining and Discovery Section, NIAMS); C. Liu (Transgenic Core, NHLBI) for their technical support. This study utilized the high-performance computational capabilities of the Biowulf Linux cluster at the NIH. This work was supported by the Intramural Research Programs of NIAMS and NIAID. H.N. was supported by the JSPS Research Fellowship for Japanese Biomedical and Behavioural Researchers at NIH.

## AUTHOR CONTRIBUTIONS

H.N. and JO’S initiated and designed the project. H.N. performed experiments, analyzed, visualized the data and drafted the manuscript. F.P. assisted H.N. in the *in situ* Hi-C. A.P. contributed the design of focused Hi-C. V.C. assisted H.N. in the computational analysis. Y.K. and JO’S supervised the project and revised the manuscript.

## DECLARATION OF INTERESTS

The authors declare no competing financial interests.

**Supplemental table 1. List of HIGs and WIGs, related to Fig. 1b**

**Supplemental table 2. List of common and cell-type specific HIGs and WIGs, related to Fig. 1c**

**Supplemental table 3. List of total and differentially regulated TADs, related to Fig. 1e**

**Supplemental table 4. gRNAs and primers for the generation of mice.**

**Supplemental table 5. Custom oligo pool for focused Hi-C**

## Methods

### Mice

All animal experiments were performed in the AAALAC-accredited animal housing facilities at NIH, by following the NIH guidelines for the use and care of live animals and were approved by the Institutional Animal Care and Use Committee of NIAMS. Mice of 6 -12 weeks old were used in all experiments. Female C57BL/6J mice purchased from the Jackson Laboratory were used for *in vitro* ILC2 and Th2 culture. The littermate mice were used for Papain-induced lung inflammation. The KHS-I KO, KHS-II KO, 5HS-I(a+b+c+d) KO, 5HS-Ie KO, 5HS-II KO and 5HS-III(a+b+c) KO strains were generated via CRISPR/Cas9 method^1^ at the transgenic core of the National Heart, Lung, and Blood Institute (NHLBI) at NIH. Briefly, a pair of single guide RNAs (sgRNAs) were designed for targeting a regulatory region, one near the 5’ end and the other near the 3’end, for the 6 predicted regulatory regions (**Table 5**). Those sgRNAs were ordered and purchased from Synthego (Menlo Park, CA). SpCas9 protein (Cat#1081059) was purchased from Integrated DNA Technologies (Corelville, IA). The pair of sgRNAs (final concentration 6ug/ml each) and Cas9 protein (final concentration 200 ug/ml) were co-electroporated into zygotes collected from C57BL/6J mice (Charles River Laboratory) using a NEPA21 electroporator (Bulldog Bio, Portsmouth, NH). Transfected embryos were cultured overnight in KSOM medium (MilliporeSigma) in a 37°C incubator with 6% CO2. In the next morning, those embryos at the 2-cell stage were implanted into the oviducts of pseudo-pregnant surrogate mothers (CD1 strain from Charles River Laboratory). Offspring were genotyped by PCR (**Table 5**), and PCR products with the predicted deletion sizes between the two sgRNA were subjected to Sanger sequencing to confirm the appropriate deletions.

### Preparation of cell suspensions from lungs and spleens

Before isolating cells from lungs, mice were perfused with 20 ml of PBS to remove blood from the tissues. The isolated lungs were placed in RPMI-1640 containing 2% (vol/vol) FCS (Thermo Fisher Scientific), 50 μg/ml Liberace TM (Roche, 05401127001), 10 μg/ml DNase I (Sigma-Aldrich, DN25) (4 ml per mouse), followed by mechanical dissociation with the gentleMACS Dissociators (Miltenyi) using gentleMACS C Tubes (Miltenyi). Samples were further digested for 45 min at 37°C. The resultant cell suspension was passed through 70 μm cell strainers, followed by additional 2 washes with PBS to recover cells. The collected single cell suspension was enriched with 40% Percoll (GE Healthcare) plus centrifugation at 400 g for 20 min, and isolated lung cells were washed twice before further analysis. For isolation of splenic cells, spleens were smashed on 70 μm cell strainers, and cell strainers were washed twice with PBS to recover cells in suspension. Splenic cells were treated with ACK buffer to remove red cells before analyses.

### Antibodies

The following antibodies were used for flow cytometry and cell purification in this study: Biotin-CD3ε (145-2C11, Biolegend), Biotin-TCRβ (H57-597, Biolegend), Biotin-CD4 (GK1.5, Biolegend), Biotin-CD8α (53-6.7, Biolegend), Biotin-CD19 (6D5, Biolegend), Biotin-CD11c (N418, Biolegend), Biotin-CD11b (M1/70, Biolegend), Biotin-TCRγδ (GL3, Biolegend), Biotin-FcεRI (MAR-1, Biolegend), Biotin-Ly6G/C (RB6-8C5, Biolegend), Biotin-NK1.1 (PK136, Biolegend), Biotin-Ter119 (TER-119, Biolegend), Biotin-EpCAM (G8.8, Biolegend), BV785-CD45 (30-F11, Biolegend), V500-CD90.2 (53-2.1, BD Bioscience), PE-CF594-CD127 (SB/199, BD Bioscience), PE-Cy7-KLRG1 (2F1, Biolegend), BUV395-GATA3 (L50-823, BD Bioscience), BUV496-CD44 (IM7, BD Bioscience), BUV615-TCRβ (H57-597, BD Bioscience), BUV737-CD4 (RM4-5, BD Bioscience), BV421-CD8α (53-6.7, Biolegend), eFlour450-Foxp3 (FJK-16s, Thermo Fisher Scientific), BV480-Siglec-F (E50-2440, BD Bioscience), BV510-IFNγ (XMG1.2, Biolegend), BV570-CD11b (M1/70, Biolegend), BV605-TCRγδ (GL3, Biolegend), BV711-CD11c (N418, Biolegend), BV750-CD90.2 (53-2.1, BD Bioscience), BV786-CD19 (1D3, BD Bioscience), AF488-IL-4 (11B11, Biolegend), PerCP-eFluor710-IL-13 (eBio13A, Thermo Fisher Scientific), PE-IL-5 (TRFK5, Biolegend), PE-CF594-KLRG1 (2F1, BD Bioscience), PE/Fire 640-MHCII (M5/114.15.2, Biolegend), PE-Cy5-CD127 (SB/199, Biolegend), PE/Fire 700-CD45 (30-F11, Biolegend), PE-Cy7-T-bet (4B10, Biolegend), AF700-NK1.1 (PK136, BD Bioscience), APC-Cy7-CD3ε (17A2, BD Bioscience), APC-Fire810-Ly6G/C (RB6-8C5, Biolegend).

### Flow cytometry

All cells were incubated with anti-CD16/CD32 (2.4G2) to block non-antigen-specific binding of immunoglobulins to the Fcγ receptors before being stained with the appropriate antibodies. Lung ILC2s were identified by lineage markers (CD3ε, TCRβ, CD4, CD8α, CD19, CD11c, CD11b, γδTCR, FcεR1, Gr1, NK1.1, Ter119) and additional markers as indicated in figure legends. For Live/Dead staining, Zombie Violet™ Fixable Viability Dye (Biolegend) was used. To detect the intracellular cytokines, cells were stimulated with PMA (50 ng/ml) and ionomycin (1 ug/ml) in the presence of Brefeldin A and Monensin for 4hr. For staining of intracellular cytokines and transcription factors, cells were fixed and permeabilized with Foxp3 staining buffer (Thermo Fisher Scientific) according to the manufacturer’s instructions. Data were acquired on a FACSCanto II (BD Biosciences) or LSR Fortessa (BD Biosciences) or Aurora (Cytek) and were analyzed with FlowJo software (Tree Star).

### In vitro ILC2 and Th2 culture

For preparation of lung ILC2s, female mice were anesthetized with isoflurane and administrated intranasally with 30 μl of IL-33 (500 ng per mouse) for consecutive 4 days to expand lung ILC2s. One day after last administration, lung cells were isolated and stained with biotin conjugated antibodies against linage markers (CD3e, TCRβ, CD4, CD8α, CD19, CD11c, CD11b, γδTCR, FcεR1, Gr1, NK1.1, Ter119, EpCAM) and anti-Biotin MicroBeads (Miltenyi Biotec #130-090-485), followed by sorting linage negative cells by MACS manual separator (Miltenyi). Lineage negative cells were further stained with fluorochrome conjugated antibodies against CD45, Thy1.2, CD127, and KLRG1 together with APC-R700 Streptavidin (BD Biosciences), followed by sorting ILC2s (CD45^+^ Lin^-^ Thy1^+^ CD127^+^ KLRG1^+^) on FACS Aria IIIu or Aria Fusion cell sorter (BD Bioscience). Sorted lung ILC2s were cultured in complete RPMI (RPMI1640 medium (Thermo Fisher Scientific) containing 10% (vol/vol) FCS, 50 μM 2-mercaptoethanol (Sigma-Aldrich), 100 U/ml penicillin and 100 μg/ml streptomycin (Thermo Fisher Scientific)) in the presence of IL-2 (100 U/ml, R&D Systems) and IL-7 (50 ng/ml, R&D systems, #407-ML). After 2 to 4 weeks, IL-2 and IL-7 were removed from ILC2 culture and additionally cultured for 4 hr before restimulation with IL-33 (Biolegend, #580508) plus NMU (R&D systems, # 1917) for mRNA-seq, ATAC-seq, ChIP-seq and HiC-seq. For *in vitro* Th2 culture, splenic naïve Th cells from mice were isolated using the Naive CD4^+^ T Cell Isolation Kit, mouse (Miltenyi Biotec # 130-104-453) according to the manufacturer’s instructions. Naive CD4+ T cells were activated by plate-bound anti-CD3 (10 μg/ml, clone: 145-2C11) and anti-CD28 (1 μg/ml, clone: 37.51) in cRPMI for 5 days with IL-2, IL-4 (50 ng/ml, R&D Systems) and anti-IFN-γ (10 μg/ml, clone: XMG1.2) for Th2 differentiation. After 5 days, cells were transferred to uncoated flasks with fresh media containing IL-2 (100 U/ml) and IL-4 (10 ng/ml). The cells were maintained at the density of 0.2∼1.2 x 10^6^ cells/ml and supplied with fresh medium every 2 or 3 days. After a total of 2 weeks in culture, cytokines were removed and cells were rested for 4 hr before re-stimulation. Th2 cells were re-stimulated with anti-CD3 and anti-CD28 for 1hr or 4hr an subjected to mRNA-seq, ATAC-seq, ChIP-seq and HiC-seq.

### PrimeFlow

mRNA expression of type 2 cytokines in ILC2s was measured by using PrimeFlow™ RNA Assay Kit (ThermoFisher Scientific # 88-18005-210) according to the manufacturer’s instructions. ILC2s were stained with Zombie Violet™ Fixable Viability Dye before fixation with RNA fixation buffer 1. The target probes against *Il4, Il13* and *Il5* mRNA were hybridized, amplified, and labeled with AF647, AF700 and AF488 respectively, followed by flow cytometry.

### Papain-mediated pulmonary inflammation in vivo

*M*ice were anesthetized with isoflurane and administrated intranasally with 30 μl of Papain (50 μg per mouse) at day 0, 3, 6 and 14 and euthanized at day 15 for analyses of lungs. One lobe of the lung was used for histological analysis. The rest of lungs were used for flow cytometry.

### Bulk mRNA sequencing

Approximately 20,000–50,000 cells (cell viability >98%) were lysed with TRIzol (Life Technologies) and total RNAs were extracted using Direct-zol™ RNA MicroPrep (Zymo Research). Total RNAs were subsequently processed to generate an mRNA-seq library using a NEBNext Poly(A) mRNA Magnetic Isolation Module (NEB, E7490S), NEBNext Ultra RNA Library Prep Kit for Illumina (NEB, E7530S) and NEBNext Multiplex Oligos for Illumina (Index Primers Set 1) (NEB, E7335S) according to the manufacturer’s instructions. The libraries were sequenced for 50 cycles (single read) with a HiSeq 3000 or NovaSeq 6000 (Illumina).

### ATAC sequencing

ATAC-seq was performed according to a published protocol^2^ with minor modification. Ten thousand cells were pelleted and washed with 50 μL 1 × PBS. After lysis, nuclei were pelleted by centrifuging at 500 × g for 10 min, and re-suspended in 50 μL transposase mixture (25 μl of 2x TD buffer, 2.5 μl of TDE1, 0.5 μl of 1% digitonin, 22 μl of nuclease-free water) (#20034197, Illumina; #G9441, Promega) for incubattion at 37°C with shaking at 300 rpm for 30 min. The tagmentated DNAs were then purified using a QIAGEN MinElute kit and amplified with 10 or 11 cycles of PCR based on the amplification curve. The libraries were purified using a QIAGEN PCR cleanup kit, and sequenced for 50 cycles (paired-end reads) on a HiSeq 3000 or NovaSeq 6000 (Illumina).

### ChIP sequencing

Stimulated or unstimulated cells were fixed for 10 min with 1% formaldehyde (ThermoFisher, #28908). Fixation was quenched by adding glycine (Sigma) at a final concentration of 125 mM. The nuclei preparation, chromatin digestion and chromatin immunoprecipitation were performed using SimpleChIP Plus Enzymatic Chromatin IP Kit (#9005, Cell Signaling Technology) with the antibodies against RNA polymerase II (#ab5408, abcam), p300 (#ab14984, abcam), GATA3 (#5852, Cell signaling), NFAT1 (#5861, Cell Signaling Technology), RelA (#8242, Cell Signaling Technology), CTCF (#07-729, Millipore Sigma) and Rad21 (#ab992, abcam). The immunoprecipitated DNAs were subsequently processed to generate a DNA library using NEBNext Ultra II DNA Library Prep Kit for Illumina (NEB, # E7645L) and NEBNext® Multiplex Oligos for Illumina (NEB, #E6440S). The libraries were sequenced for 50 cycles (single read) with a HiSeq 3000 or NovaSeq 6000 (Illumina).

### Global and focused HiC sequencing

A detailed protocol to generate HiC libraries can be obtained at PMID:25497547 and PMID:26499245^3, 4^. For focused HiC-seq, we designed 1490 probes targeting chr11:53,250,000 – 54,250,000 (length: 1Mb) containing type 2 cytokine locus (**Table 4**). The libraries were sequenced for 50 or 100 cycles (paired-end read) with a HiSeq 3000 or NovaSeq 6000 (Illumina).

### Histological analysis

Lungs were fixed for at least 24 h with 10% formalin (Sigma-Aldrich, #HT5012) and were embedded in paraffin. Cross-sectional lung tissues were stained with Periodic acid–Schiff (PAS) or hematoxylin and eosin. Pathological score was evaluated by an observer masked to treatment group for the following parameters: peribronchiolar inflammation and cuffing (0–4), perivascular inflammation and cuffing (0–4), goblet cell hyperplasia (0–4) and interstitial infiltrate (0–3).

### Bulk mRNA-seq analysis

Raw sequencing data were processed with CASAVA 1.8.2 (Illumina)^5^ to generate FastQ files. Sequence reads were mapped onto mouse genome build mm10 using TopHat 2.1.0^6^. Gene expression values (FPKM; fragments per kilobase exon per million mapped reads) were calculated with Cufflinks 2.2.1^6^. BigWig tracks were generated from Bam files and converted into bedGraph format using bedtools^7^. These were further reformatted with the UCSC tool bedGraphToBigWig and visualized by Igv 2.11.9^8^. Published mRNA-Seq data from ILC2s (GSE131996)^9^ were included in the final analysis. Downstream analyses and graph generation were performed with R 4.1.1^10^.

### ATAC-seq analysis

ATAC-seq was done in two biological replicates per condition. Reads were mapped to the mouse genome (mm10 assembly) using Bowtie 0.12.8^11^. In all cases, redundant reads were removed using FastUniq^12^, and customized Python scripts were used to calculate the fragment length of each pair of uniquely mapped paired-end (PE) reads. Distributions of fragment sizes were similar to previously published data (data not shown). Reads whose fragment lengths were less than 175 bp were kept and only one mapped read per a unique genomic region was used to call peaks. Regions of open chromatin were identified by MACS (version 1.4.2)^13^ using a p-value threshold of 1 × 10^−5^. Only regions called in both replicates were used for downstream analysis. Peak annotation and motif analysis were performed with the Hypergeometric Optimization of Motif EnRichment program (HOMER) version 4.11.1^14^ using the following parameter settings; “annotatePeaks.pl peak_file mm10 -size 1000 -hist 40 -ghist” and “findMotifsGenome.pl peak_file mm10 motif_folder -size given -preparsedDir tmp 2 > out”. Published ATAC-Seq data from ILC2s (GSE131996)^9^ and from human Th cells (GSE126117)^15^ were included in the final analysis.

### ChIP-seq analysis

Sequencing data from ChIP-Seq were mapped onto mouse genome build mm10 using Bowtie 0.12.8^11^. All ChIP-Seq experiments were done in biological duplicates. BigWig tracks were generated from Bam files and converted into bedGraph format using bedtools^7^. These were further reformatted with the UCSC tool bedGraphToBigWig. Tag directory were generated by the HOMER 4.11.1^14^ and visualized by Igv 2.11.9^8^. MACS (version 1.4.2)^13^ was used for peak calling using a P value threshold of 1 × 10−5. Quantification of p300 load and RelA load in the +/- 50kb from TSS of HIGs was performed with HOMER as follows; “annotatePeaks.pl tss mm10 -size 100000 -list list.HIGs.txt -d ChIP.tagdir > output.txt”. Downstream analyses and graph generation were performed with R 4.1.1^10^.

### HiC-seq analysis

All HiC-Seq experiments were done in biological duplicates. The initial Hi-C data processing was done with HOMER 4.11.1^14^. The MboI restriction sites used for self ligation (GATC) were trimmed from the FastQ files using “homerTools trim” function. The trimmed reads were mapped onto mm10 using Bowtie 2.4.5^11^. The paired-end tag directories were created using “makeTagDirectory” command, which also generated several quality control analysis files. The .hic files were created using HOMER function “tagDir2hicFile.pl” with Juicer 1.6^16^ and visualized with Juicebox 1.9.8. TADs and Loops were detected with HOMER using “findTADsAndLoops.pl find tagdir_file -cpu 10 -res 1000 -window 5000 -genome mm10”. This program also generated a bedGraph that describes the directionality index (DI). To identify differentially regulated TADs, TAD files from all experiments were merged using “merge2Dbed.pl” command followed by quantifying TADs across samples using “findTADsAndLoops.pl score -tad merged.tad.2D.bed -d hic.tagdirs -cpus 10 -o output”. The circos plots showing the chromatin interaction within *Il4/Il13/Il5* (chr11:53,400,000-53,800,000), *Il3/Csf2* (chr11:54,100,000-54,600,000) and *Il9* locus (chr13:56,400,000-56,900,000) with ChIP-seq data were generated with HOMER function using “analyzeHiC hic.tagdir -pos region -res 1000 -superRes 2000 -circos filename -nomatrix -d CTCF.tagdir/ Polll.tagdir/ p300.tagdir/ GATA3.tagdir/ RelA.tagdir/ NFAT1.tagdir/ -g mm10.genes -pvalue 0.0001 -minDist 10000”. The normalized HiC contact matrices with 1kb bins were obtained by HOMER function “analyzeHiC hic.tagdir -pos region -res 1000 -window 2000 -balance -override > hic.matrix.txt”. The matrix files were reformatted with HiCExplorer 3.6^17–19^ using the command “hicConvertFormat -m hic.matrix.txt --inputFormat homer --outputFormat h5 -o hic.matrix.h5” and created heatmap with HiCExplorer using “hicPlotsTADs”. Published HiC-Seq data from human Th cells (GSE126117)^15^ were included in the final analysis. Further downstream analyses and graph generation were performed with R 4.1.1^10^ and Prism 8.

### 3D modeling of type 2 cytokine locus

The normalized HiC contact matrices of type 2 cytokine locus (chr11:53,425,000-53,775,000, 350kb) with 1kb bins were generated with HOMER as an input for 3D modeling with GenomeFlow v2.0^20^. The LorDG function was used to reconstruct 3D chromatin using Conversion Factor of 0.5, Initial Learning Rate of 1.0 and Max Number of Interactions of 1000. The distance between bins containing CGRE (chr11: 53635000-53635999), RHS(5+6) (53654000-53654999), 5HSIs (53713000-53713999), 5HSII (53731000-53731999), 5HSIIIs (53734000-53734999), +6.5kb*^Il^*^13^ (53627000-53627999), HSII (53616000-53616999), KHSI (53584000-53584999) was calculated by LorDG function and visualized with R 4.1.1^10^.

### Statistics and reproducibility

All experiments shown in this work were performed at least twice with consistent results. The presented data were collected from biologically independent samples. Sample size was not predetermined. Unless otherwise indicated, data are presented as the mean with standard error (s.e.m.) to indicate the variation within each experiment. Statistical analyses were performed in Excel, Prism and R. Unless otherwise indicated, a two-tailed t-test was used for assessing significance.

## DATA AND SOFTWARE AVAILABILITY

The data of mRNA-seq, ATAC-seq, ChIP-seq and Hi-C generated in this study will be deposited in the NCBI Gene Expression Omnibus (GEO) before publication.

---

**Extended Data Figure 1.**
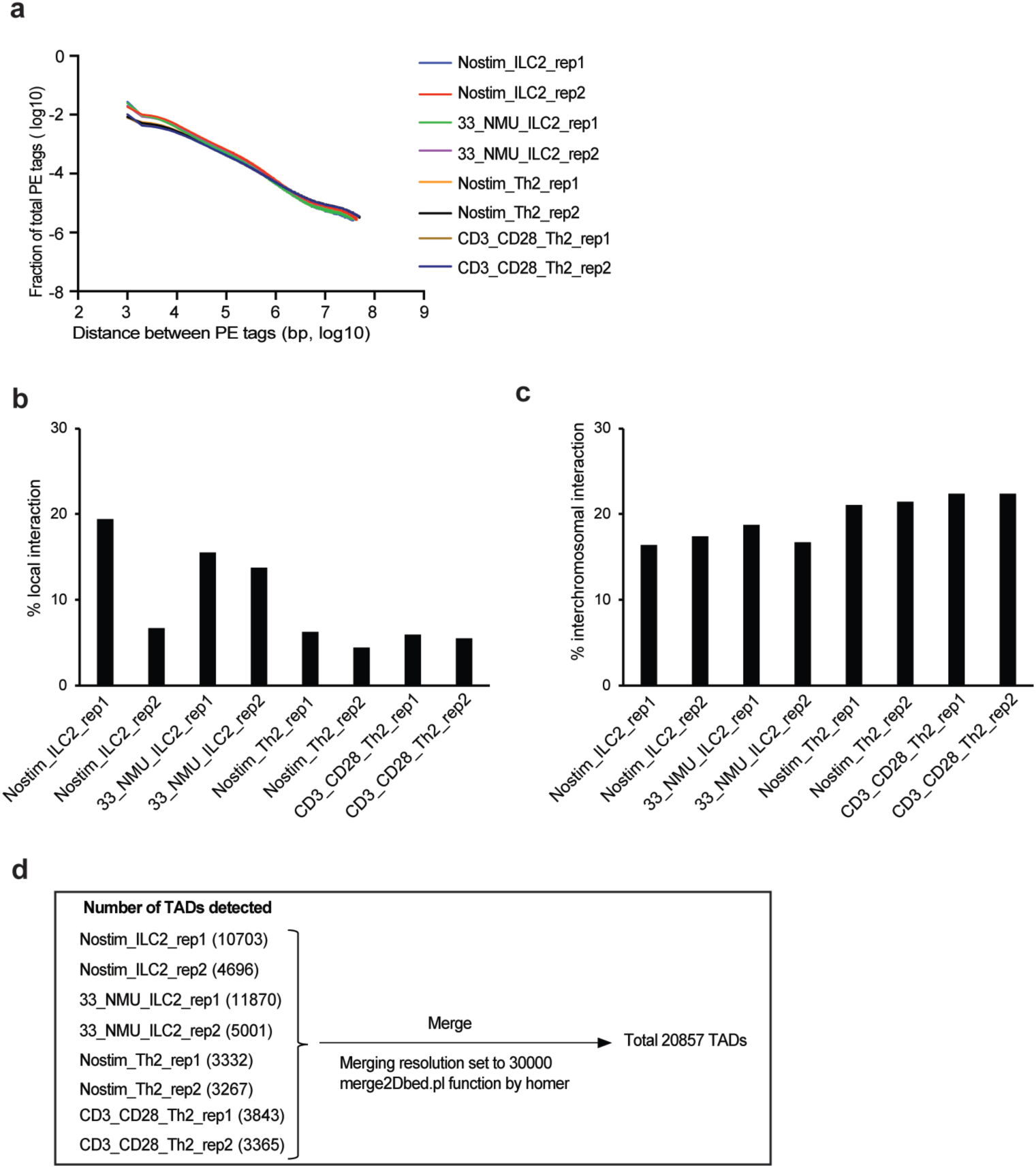
in situ Hi-C of ILC2s and Th2 cells. **a,** Interaction frequency vs. distance. **b,** Percentage of local interactions. **c,** Percentage of interchromosomal interactions. **d,** Number of TADs detected in individual samples. All TADs are merged by merge2Dbed.pl function by homer for downstream analysis.

**Extended Data Figure 2.**
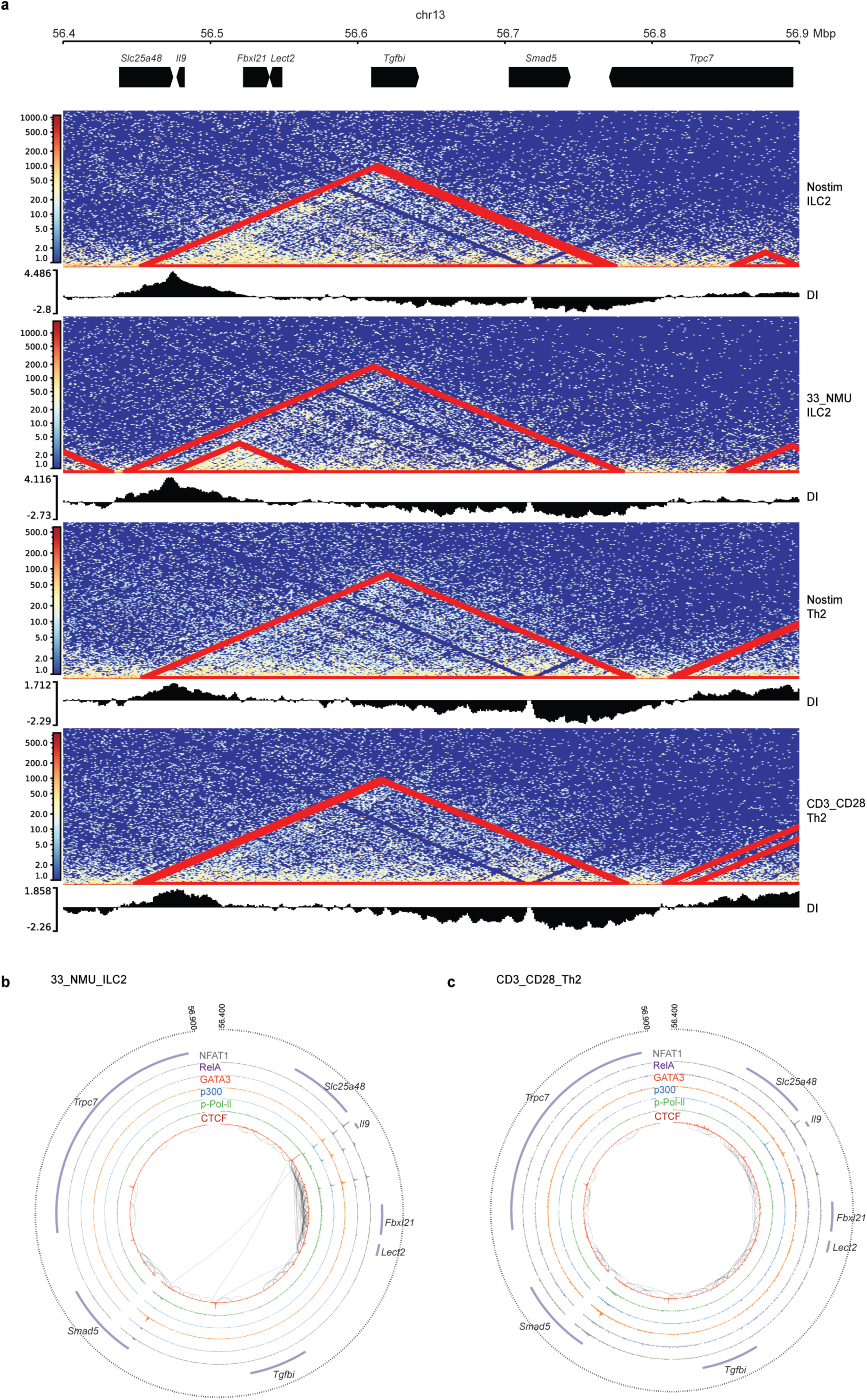
Chromatin remodeling at Il9 locus. **a**, HiC heatmap at 56.4 – 56.9 Mbp on chromosome 13 containing *Il9* locus. DI: directionality index. **b, c**, Circus plots representing HiC interactions and TF footprints at 56.4 – 56.9 Mbp on chromosome 13 in ILC2s (**b**) and Th2 (**c**) following activation.

**Extended Data Figure 3.**
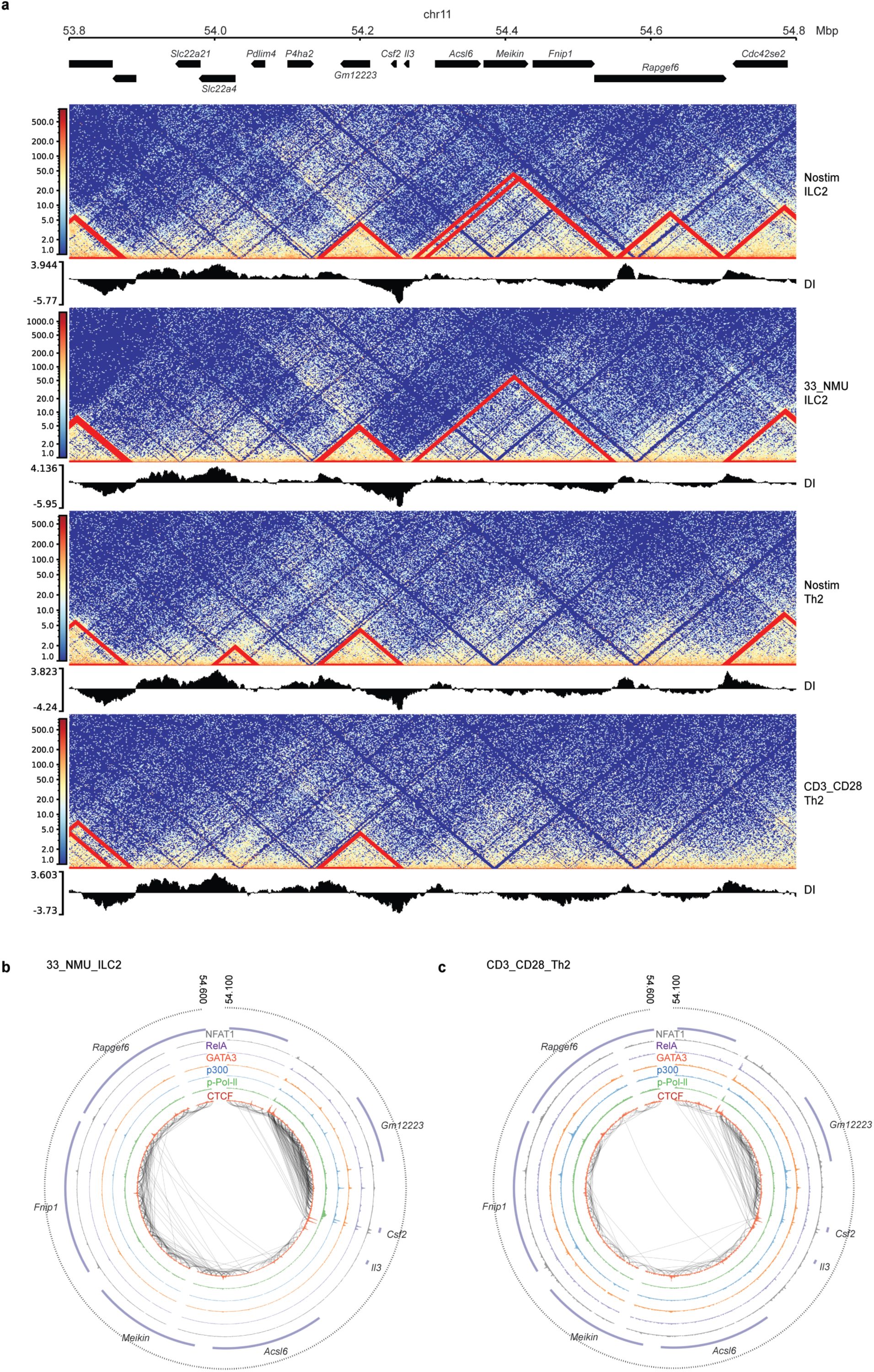
Chromatin remodeling at Csf2 / Il3 locus. **a**, HiC heatmap at 53.8 – 54.8 Mbp on chromosome 11 containing *Csf2 and Il3* locus. DI: directionality index. **b, c**, Circus plots representing HiC interactions and TF footprints at 53.8 – 54.8 Mbp on chromosome 11 in ILC2s (**b**) and Th2 (**c**) following activation.

**Extended Data Figure 4.**
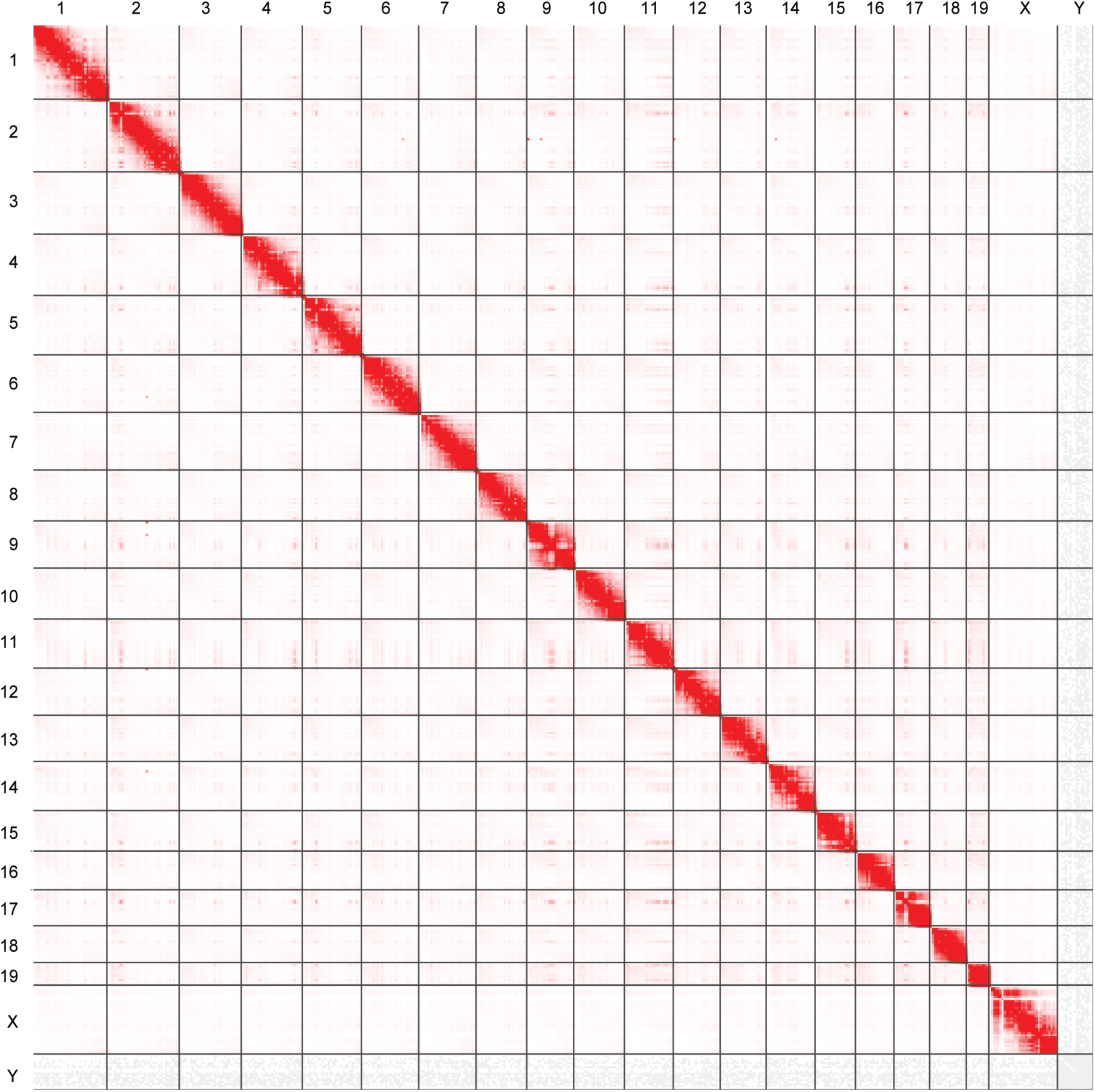
Overview of intra- and inter-chromosomal interaction in ILC2s. Genome-wide view of intra- and inter-chromosomal interactions in stimulated ILC2s, analyzed by Juicebox.

**Extended Data Figure 5.**
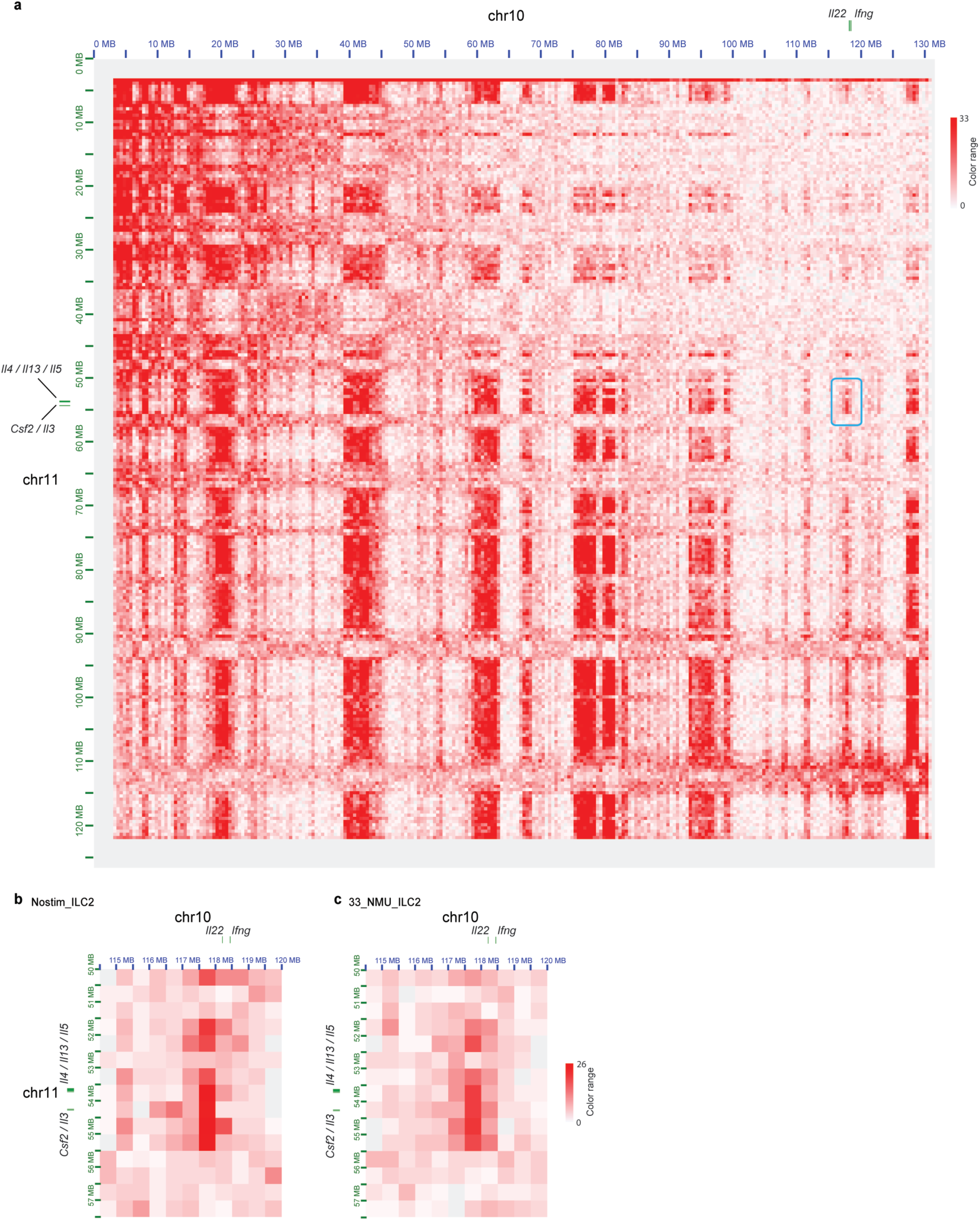
Interchromosomal interaction between chr 10 and chr 11 in ILC2s. **a**, Overview of interchromosomal interaction between chr 10 and chr 11 in non-stimulated ILC2s. *Il22 / Ifng* on chr 10 and *Il4 / Il13 / Il5 / Csf2 / Il3* on chr11 were depicted respectively. **b, c**, Extended interaction between the regions containing *Il22 / Ifng* loci and *Il4 / Il13 / Il5 / Csf2 / Il3* loci in ILC2s during homeostasis (**b**) and following activation (**c**).

**Extended Data Figure 6.**
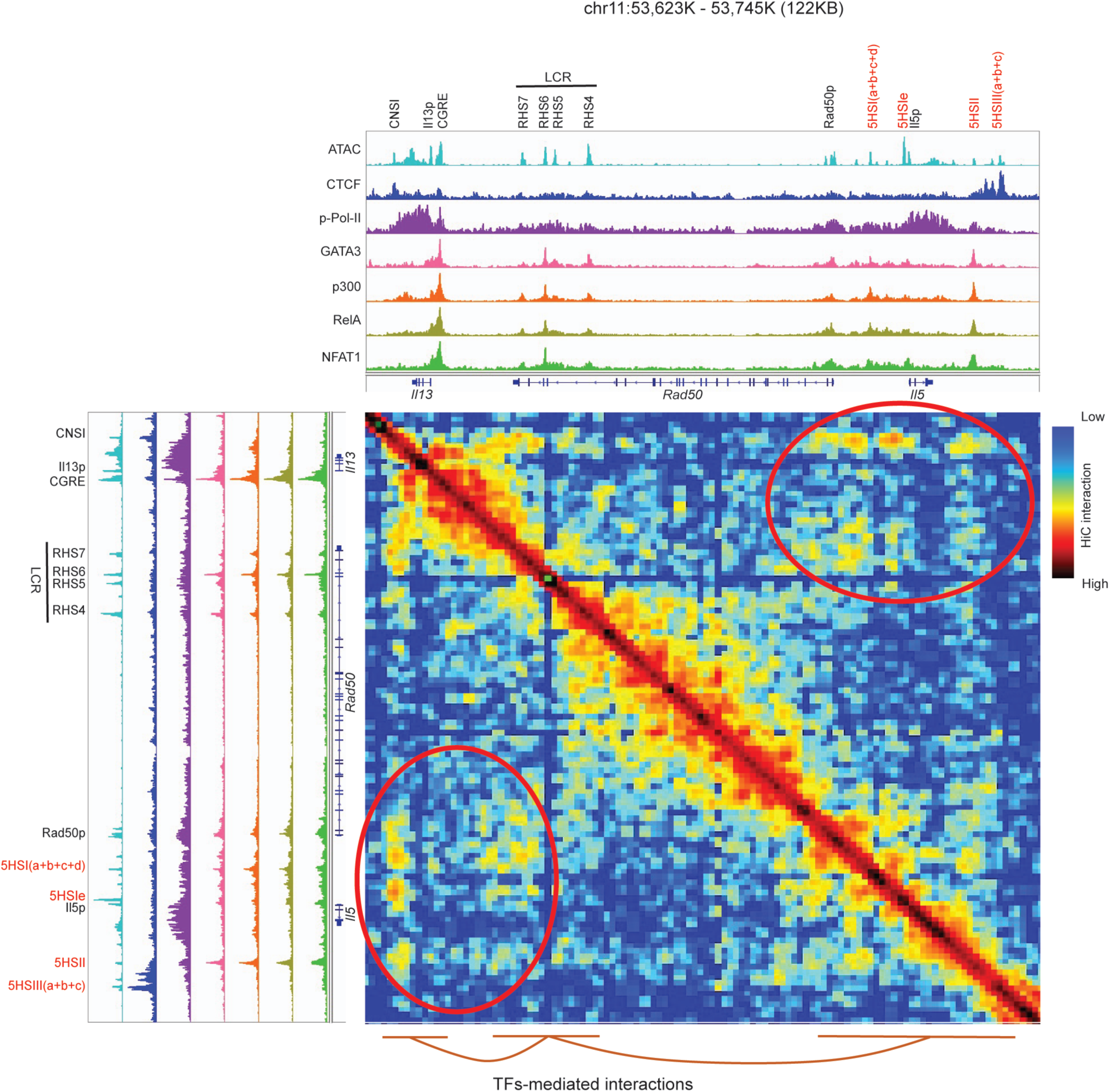
Remodeling of Il13/Rad50/Il5 loci induced by SDTFs. A compiled view of *Il13/Rad50/Il5* loci showing chromatin accessibility, TFs binding and HiC interaction in ILC2s stimulated with IL-33 plus NMU. Data represents chr11: 53,623,000-53,745,000 (122kB).

## Notes

### Competing Interest Statement

The authors have declared no competing interest.

